# TCR clonality and TCR clonal expansion in the *in situ* microenvironment of non-small cell lung cancer

**DOI:** 10.1101/2024.12.17.628857

**Authors:** Hui Yu, Anastasia Magoulopoulou, Rose-Marie Amini, Maria Paraskevi Chatzinikolaou, Masafumi Horie, Amanda Lindberg, Artur Mezheyeuski, Max Backman, Andreas Metousis, Hans Brunström, Millaray Marincevic Zuniga, Johan Botling, Johanna Sofia Margareta Mattsson, Klas Kärre, Karin Leandersson, Mats Nilsson, Carina Strell, Patrick Micke

## Abstract

T-cell activation and clonal expansion are essential for the efficacy of immunotherapy in non-small cell lung cancer (NSCLC) patients. Since the distribution of T-cell clones might provide insights into immunogenic mechanisms, we determined the α/β TCR clonality using RNA-sequencing from frozen tumor tissue of 182 NSCLC patients and paired the results with extensive *in situ* image and sequence analyses of the immune microenvironment of NSCLC. TCR clonality (Gini index) patterns ranged from high T-cell clone diversity with high evenness (Gini index low) to clonal dominance with low evenness (Gini index high). TCR clonality in cancer tissue was lower than in matched normal lung (p=0.021). High Gini index correlated strongly with distinct mutations (EGFR, P53), tumor mutation burden (p<0.001), and inflamed tumor phenotypes (PRF1, GZMA, GZMB, INFG) with exhaustion signatures (LAG3, TIGIT, IDO1, PD-1, PD-L1). Correspondingly, PD-1+, CD3+, CD8A+, CD163+, and CD138+ immune cells infiltrated cancer tissue with high TCR clonality. *In situ* sequencing revealed that dominant T-cell clones were more often of CD8-subtype and tended to approximate the tumor cell compartment (p<0.03). In a checkpoint inhibitor-treated NSCLC patient cohort, high TCR clonality was associated with therapy response (p=0.016) and prolonged survival (p=0.003, median survival 13.8 vs 2.9 months).

Our robust analysis pipeline revealed diverse TCR repertoires related to genotypes and immune phenotypes. The *in situ* positioning of expanded T-cell clones indicated functional impact, which was clinically confirmed in NSCLC patients receiving immunotherapy.

**One sentence summary:** T-cell clone expansion in NSCLC is associated with genetic mutations, immune phenotypes, immunotherapy response, and patient survival.

## Introduction

The introduction of new immune modulatory treatment options has revolutionized clinical oncology and provided clinical evidence that the immune system is able to recognize and eliminate cancer cells. In line with this, the infiltration of immune cells in the tumor tissue is associated with improved prognosis independent of therapy(*1, 2*). This includes densities of T-cell subsets (CD4, CD8, T-reg cells, CD45 T memory cells), B-cells, plasma cells, as well as subsets of macrophages and dendritic cells (*3–5*). A particular Immune cell infiltration is often associated with higher expression of T-cell activation markers but also of T-cell exhaustion (*3, 6, 7*). This immune cell-dense tumor tissue is frequently denoted as “immune hot” or “inflamed” and is not only prognostic but has also proven to be predictive in the context of immune checkpoint blockade (*8, 9, 4, 10, 11*). However, the prognostic and predictive value is still relatively low, which suggests that these quantitative measurements do not entirely reflect the patient individual actual immune reaction.

The efficacy of a specific tumor-antigen T-cell response is reflected by the precedent activation, differentiation, and clonal expansion of naive T-cells upon specific binding of their TCRs to tumor-antigens presented on MHC, together with costimulatory signals (e.g. CD28) (*12, 13*). The activation is reinforced by autocrine or paracrine IL2 signaling, contributing to massive clonal expansion of tumor-antigen specific T-cell clones (*14, 15*) generating both effector and memory T-cells (*16*). This T-cell activation, with subsequent expansion, can be measured by the clonal unevenness of the TCR in the T-cell population, which is often referred to as high clonality (*17, 18*).

TCR clonality was previously determined by targeted sequencing of the TCR-ß chain (*19*). Clinically certified assays are used to determine malignant clones in hematological malignancies that can be followed over time (*20*). In solid tumors, target sequencing methods were mostly done on blood samples to evaluate specific anti-tumor T-cell responses and their relation to clinical outcome (*21, 22*). In melanoma, tumor TCR clonality was associated with benefit from checkpoint inhibitor treatment in several studies (*23–26*).

Recently, bioinformatic algorithms have been developed to determine the clonality in crude RNAseq data without the necessity of specific assays (*27, 28*). Such RNAseq-based clonality analysis of lung cancer tissue indicated that the TCR richness, i.e., the number of different T-cell clones, was lower in the tumor tissue than in the adjacent normal tissue and was associated with histomorphological tumor features (*29*). However, T-cell repertoire analysis was most often limited to the in silico processing of public data sets, and consequently, the pure quantification of the T-cell clones excluded the cellular tissue context. It can be speculated that the TCR clonality and differentiation also depend on molecular features of tumor and stroma cells and that the individual immune microenvironment is a cause or consequence of a specific T-cell expansion. This was recently demonstrated in an elegant study characterizing “supportive” and “non-supportive” CD8 T cell niches in different cancer type (*30*).

With this background, our study aimed to determine the T-cell repertoires in the immune microenvironmental context using a thoroughly validated TCR sequencing pipeline (*31*), providing the clinical and molecular background of T-cell clonality in cancer tissue from non-small cell lung cancer (NSCLC) patients.

## Materials and Methods

### Patient material

This study includes 182 NSCLC patients from the Uppsala II cohort (n=357) for which RNAseq data were available, as described before (*32*). Patient characteristics are listed in **Table 1**. All patients from the Uppsala II cohort were NSCLC patients treated surgically at Uppsala University Hospital between 2006 and 2010. This study was conducted in concordance with the Declaration of Helsinki and the Swedish Ethical Review Act (approved by the Ethical Review Board in Uppsala, #2012/532).

**Table 1:** Patient characteristics of the NSCLC patient cohort. Patients were operated at the Uppsala University Hospital (Sweden) between 2006 and 2010. Fresh tissue was procured freshly after the operation and used later for RNAseq.

|  | No. | % |
| --- | --- | --- |
| **All patients** | **182** | **100%** |
| **Age(years)** |  |  |
| >70 | 73 | 40% |
| ≤70 | 109 | 60% |
| **Sex** |  |  |
| Female | 95 | 52% |
| Male | 87 | 48% |
| **Smoking** |  |  |
| Ever smokers | 165 | 91% |
| Never smokers | 17 | 9% |
| **Histology** |  |  |
| Adenocarcinoma | 107 | 59% |
| Squamous cell carcinoma | 62 | 34% |
| other types | 13 | 7% |
| **Stage (8<sup>th</sup> edition)** |  |  |
| 1A2 | 39 | 21% |
| 1A3 | 35 | 19% |
| 1B | 23 | 13% |
| 2A | 14 | 8% |
| 2B | 30 | 17% |
| 3A | 35 | 19% |
| 3B | 6 | 3 % |
| **Performance status (WHO)** |  |  |
| 0 | 114 | 63% |
| 1 | 66 | 37% |
| 2 | 2 | 1% |

RNAseq data from publicly available NSCLC cohort downloaded from Gene Expression Omnibus (GSE126044) were also used. These RNAseq data are derived from 9 fresh frozen samples from anti-PD1 (nivolumab) treated NSCLC patients (8 male, one female; 3 adenocarcinomas, 6 squamous cell cancer, median age 65 year) (*33*).

### RNA sequencing

Transcriptomic profiling was performed on fresh frozen tumor tissue, as described previously (*32*). RNA was extracted using RNeasy Mini Kit (Qiagen) from archived fresh frozen tumor tissues stored at -80°C. The samples were prepared for sequencing using the Illumina TruSeq RNA Sample Prep Kit v2 and poly-A selection. One hundred base paired-end multiplex sequencing was performed on the Illumina HiSeq2500 machine (Illumina, Inc. USA) with five samples per lane following the standard Illumina RNAseq protocol. The raw data is accessible in the GEO repository (accession number GSE81089).

### TCR clonality analysis

TCR clones within each sample were extracted from bulk RNAseq results using MiXCR version 3.0.13 following the software reference document (https://docs.milaboratories.com/) on a Windows 10 system sever (Java version 17.0.2, processer: Intel i9-11900). Raw RNA sequencing data was aligned with the command “mixcr align -s hs -p rna-seq - OallowPartialAlignments=true forward_sequence reverse_sequence align_output.vdjca”. After that, two rounds of “assemblePartial” were performed according to the software reference document with commands “mixcr assemblePartial align_output.vdjca assemble1.vdjca” and “mixcr assemblePartial assemble1.vdjca assemble2.vdjca”. The extension step was then performed using the command “mixcr extend assemble2.vdjca extend.vdjca”. Clones were assembled using “mixcr assemble extend.vdjca clones.clns”. TRA and TRB clones were exported separately using the commands “mixcr exportClones -cloneId -count -fraction -vGene -dGene -jGene -cGene -vAlignment -jAlignment -targets -nFeature CDR3 -c TRA clones.clns tra.txt” and “mixcr exportClones -cloneId -count -fraction -vGene -dGene -jGene -cGene - vAlignment -jAlignment -targets -nFeature CDR3 -c TRA clones.clns tra.txt”. TRA and TRB clones for each sample were merged, and a fraction of each clone was recalculated using a Python script. Clonality was defined using the Gini coefficient (Gini index)(*17, 18*) and calculated with a Python script. All the Python scripts are available as supplementary material and have been deposited on GitHub. The Gini index, the total number of clones, and clinical data for each patient are provided in **table S1**. A detailed clonal profile for each patient is provided upon request (HY).

### Differential gene expression analysis

The differential gene expression analysis was performed using the “DESeq2” package (version 1.40.2) in R. Two groups were compared: the top quartile against the bottom quartile of patients ranked by Gini index. The gene expression raw count data and the patient group records were formed as required by the software instruction. “DESeqDataSetFromMatrix”, “DESeq”, and “results” from the “DESeq2” package was used with the default function. The full result is shown in the **table S2**. The GO and KEGG analysis was performed using “enrichGO” and “enrichKEGG” with default parameters from package “clusterProfiler” (version 4.8.2).

### Lymphotrack

Four patients with different clonality/Gini index were analyzed by the Lymphotrack assay (*20*). Three 10 µm fresh frozen tissue sections of each patient sample were transferred into ATL buffer, and DNA was extracted using DNeasy Blood & Tissue kit (Qiagen Germany) following the manufacturer’s protocol. Extracted DNA was stored in 100 µL AE buffer, and the concentration was measured by the Qubit 2.0 Fluorometer (Invitrogen) using QubitTM dsDNA BR Assay Kit (Invitrogen, USA) according to the manufacturer’s protocol. TRB and TRG clones were measured respectively using LymphoTrack Dx TRB Assay Kit A-MiSeq (Invivoscribe, Inc. USA) and LymphoTrack Dx TRG Assay Kit Panel-MiSeq (Invivoscribe) on the Illumina MiSeq sequencing platform. Raw sequence data was analyzed and visualized using the LymphoTrack Software-MiSeq (version 2.4.3, Invivoscribe).

### Immunohistochemistry

Immunohistochemistry was conducted as described previously (*32, 34*), and the detailed protocol is available on the human protein atlas website (www.proteinatlas.org/download/IHC_protocol.pdf). Formalin-fixed, paraffin-embedded (FFPE) tissue microarrays (TMA), including two 1mm (diameter) representative punches of each patient, were cut to 4 µm sections. Automated IHC was conducted using the Autostainer 480 (Thermo Fisher Scientific, USA) as described before (*9, 32*), The antibodies used were: CD3 (CL1497, 1:1000 dilution, Atlas Antibodies), CD4 (CL0395, 1:125 dilution, Atlas Antibodies), CD8A (CL1529, 1:250, Atlas Antibodies), CD20 (L26, pre-prepared manufacturer dilution; Agilent Technologies, USA), CD45RO (UCHL1, 1:1000 dilution; Abcam, UK), CD138 (MI15, 1:100 dilution; Agilent Technologies), CD163 (10D6, 1:100 dilution; Novocastra, Leica Biosystems, USA), FOXP3 (236A/E7, 1:15 dilution; Santa Cruz Biotechnology, USA), PD1 (MRQ-22, 1:100 dilution; Cell Marque, USA), and NKp46 (195314, 1:50 dilution; R&D Systems, USA). PD-L1 (22C3, ready-to-use solution, Agilent Technologies) staining was performed at the Clinical Pathology Unit at Uppsala University Hospital on a DAKO autostainer system according to manufacturer instructions.

Afterward, vision-based manual annotation was carried out to calculate the immune score on the stroma and tumor area of each TMA core, as described before (*9, 35, 36*). The immune score was calculated by dividing stained, positive immune cells by all other cells in the stroma or tumor compartment respectively. The PD-L1 staining annotation in the tumor compartment followed the standard annotation of cancer cell staining used in clinical diagnostics, in which partial or complete membrane-stained tumor cells were all counted positive.

### Multiplex immunofluorescence

Multiplex immunofluorescence staining data were previously described (*4, 36*). In brief, TMA sections were stained with antibody panels: a tumour-infiltrating lymphocyte panel (TIL), including CD4, CD8, CD20, FoxP3 and PanCK; a natural killer cell/macrophage panel (NK/MP), including CD3, NKp46, CD56, CD68, CD163 and PanCK; and an antibody presenting cell panel (APC), including CD1a, CD208, CD123, CD15, CD68 and PanCK. The staining procedure was performed in accordance with an earlier published study (*35, 36*) based on the modified Opal Multiplex IHC assay (Akoya Biosciences, USA). The slides were scanned using the VectraPolaris system (Akoya Biosciences) at a 2 pixels/μm resolution in multispectral mode and analysed in the inForm software, where spectral unmixing was used to generate an oligo-layer image with layers corresponding to the specific staining, 4′,6- diamidino-2-phenylindole (DAPI), and autofluorescence. The inForm software was used to define tumor and stroma compartments within each tissue core. The algorithm was trained on pathologist-annotated samples. Cell segmentation was based on DAPI nuclear staining. Representative subsets of the included markers were annotated as either positive or negative, and the inForm software was then trained on these to phenotype all other cells accordingly. The intensity of each marker expression was used to calculate the thresholds for marker positivity. A pathologist reviewed each image and curated it with regard to artefacts, staining defects and necrosis.

### Mutation analysis

As described previously (*37*), genomic DNA was extracted from fresh frozen tissue, using the QIAamp DNA Mini Kit (Qiagen) or the QIAamp DNA FFPE Tissue (Qiagen) respectively. Targeted DNA enrichment was performed using the Haloplex target enrichment system (Agilent Technologies), and targeted deep sequencing was conducted on the Illumina HiSeq 2500 platform following the manufacture protocol. La Fleur et al. performed downstream analysis and identified gene variants as published before.

### *In situ* sequencing

The sequencing was performed with modification as previously described (*38, 39*). Tumor sections (10 µm) from frozen tumor tissue slides were thawed and air-dried at room temperature for a maximum of 5 minutes. Fixation was performed with 3.7% formaldehyde (FA) for 10 minutes and the samples were rinsed twice in PBS. Permeabilization followed immediately, using 0.1 M Hydrochloric Acid (HCl) diluted in water, at room temperature for 5 minutes. Two PBS washes followed, each lasting 1 minute, and then dehydration with 70% and 100% ethanol for 2 minutes each. After air drying, SecureSeal Hybridization chambers by Grace Bio-Labs were mounted on the tissue sections.

The padlock probe hybridization and rolling circle amplification (RCA) were performed according to the High Sensitivity Library Preparation Kit by Cartana (Cartana, 10x Genomics, USA). In brief, the padlock probe mix (including the Immune General P/N: 4121-13 Lot: 4WW44632, Immune Oncology P/N:4122-13 Lot:4QZ44362, and TCR padlock probe panels together with blocking probes for the high expressors IGKC, PTPRC, and LST1, Supplementary table 6-7) was diluted in Buffer A, and incubated at 37 °C overnight. On the second day of the protocol, a 30-minute wash with WB4 at room temperature and a 2-hour ligation with RM2 and Enzymes 1 & 2 at 37°C was performed. The RCA step was then carried out overnight at 30 °C with RM3 and enzyme 3 mix.

The procedure continued with the hybridization of L-probes and detection probes on the RCPs, using Cartana’s In Situ Sequencing Kit (Cartana). Initially, the samples were incubated with the L-probe mix for an hour at 37°C and washed thrice with PBS. Next, they were incubated with the detection oligo mix for an hour at 37°C and again washed thrice with PBS. Autofluorescence was reduced with Vector® TrueView® from Vector Laboratories (Vector Laboratories, Inc. USA), as per the manufacturer’sprotocol. Cover slips were mounted with SlowFade™ Gold antifade reagent by Invitrogen (Invitrogen Crop, USA). Lastly, the protocol included cover slip removal and detection oligo stripping between imaging rounds, with 100% formamide for three one-minute incubations, as outlined by Cartana’s In Situ Sequencing Kit. Image acquisition and data processing were performed as described previously.(*38*) The entire code is available at https://github.com/Moldia/Lee_2023. The probes were listed in **table S6-S7**.

### Hexbin based *in situ* sequencing data analysis

The analysis of the sequencing result was performed with Python code (version 3.10.13, https://github.com/JnuYHui/TCR_in_NSCLC). The DAPI image from sequencing was segmented by hexagon bins (long diagonal 300 µm). The sequencing signals were allocated to each hexbin according to their coordinates. The HE staining was performed on the same slide after imaging for DAPI staining according to the standard HE staining protocol. The HE staining slides were scanned using bright field scanning and overlayed to the DAPI image using QuPath (0.5.1) and Python scripts. The hexbins with either EPCAM or CDH1 expression (cutoff = 2 signal counts) were assigned as tumor compartment. The empty or necrotic areas were manually annotated. The remaining hexbins were assigned as stroma compartments. TCR clones were considered as dominant with a count larger than 10 in the bulk RNAseq data. TCR clones were recovered in the *in situ* sequencing data using Python scripts, and the number of dominant clones were counted in each tumor and stroma hexbin and differences were assessed by chi-square test. Dot plots were made using Python scripts and illustrate the fraction and expression distribution of TCR clones. The stack plots were made in OriginPro2024b (OriginLab Corporation, Massachusetts, USA).

### Statistics and bioinformatics

Data analysis was performed using R (version 4.2.0). Median TCR clonality (Gini index = 0.25431) was used as a cut-off to define high and low clonality. Statistical significance was set to p < 0.05. Unpaired Wilcoxon rank sum test was used to compare two independent groups, while the unpaired Kruskal-Wallis test was used for more than two independent groups. Boxing plots and the significant test were produced with the ggplot2 package (version 3.3.6). Overall survival analysis was performed using the Kaplan-Meier method with R package “survival” (version 3.3-1) and “survminer” (version 0.4.9) using the Log-Rank test for group comparison. Multivariate Cox regression test from the “survival” package was used to calculate the relative hazard ratios (HR) with 95% confidence intervals (CI) controlling for all causes. The best cut- off calculation was performed using the “surv_cutpoint” function from the “survminer” package. Spearman’s rank correlation between Gini index and the immune markers was calculated using the Hmisc package (version 4.7-1). Hierarchical cluster analysis was carried out using “ward.D2” as a clustering method measuring the Euclidean distance using the “pheatmap” package (version 1.0.12). The DESeq2 package (version 1.36.0) was used to perform differential gene expression. GO and KEGG analysis of the significantly differentially expressed genes were performed using the clusterProfiler package (version 4.4.4). Chi-square tests were used comparing fraction of dominant TCR clones in tumor and stroma hexabins (Python). The Python scripts were deposited at GitHub (version 3.10.13, https://github.com/JnuYHui/TCR_in_NSCLC).

## Results

### The T-cell repertoire in lung cancer

The T-cell repertoire was determined on RNAseq data derived from 182 tumor tissues from operated NSCLC patients (**Table 1**). In addition, 20 paired normal lung tissues distant from the tumor of the same patients were analyzed. Quantifying RNA reads attributed to each T-cell receptor determines the relative frequency and proportion of TCR clones in each sample. These proportions are then visualized using pie charts. (**Fig. 1a**). To transfer the pattern of T-cell clones to a metric, we applied the Gini coefficient (Gini index) (*17*). A T-cell repertoire with few dominant clones results in a high Gini index, whereas a repertoire without dominant clones, with even distribution, results in a low Gini index (**Fig. 1a**). Using the DNA-based LymphoTrack assay (*20*), we recovered the identified dominant clones in four selected samples (**Fig. S1**). The agreement indicates the validity and robustness of our method and the quality of the RNA data sets.

**Fig. 1.**
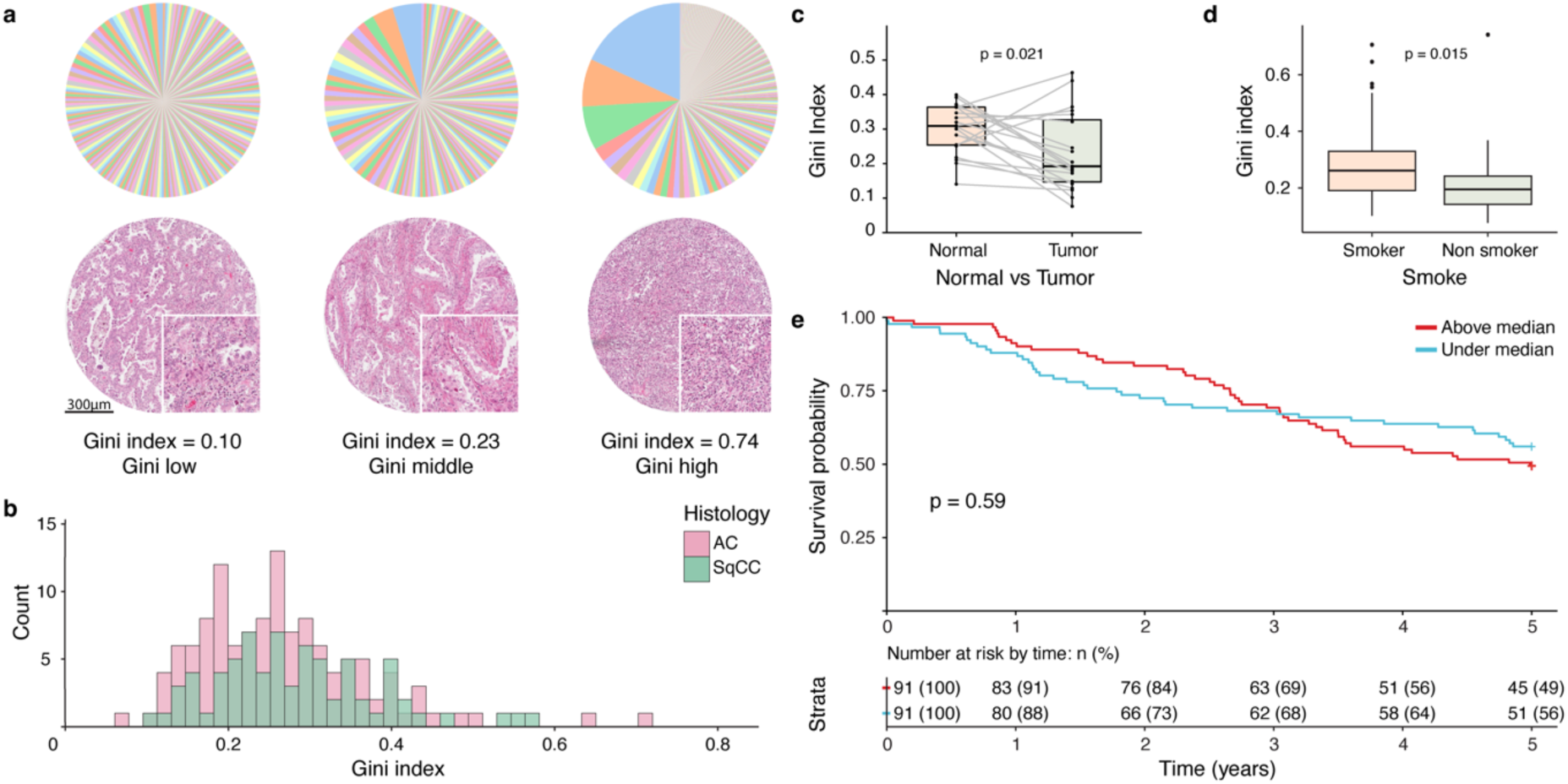
TCR clonality in tumors of NSCLC patients. **(a)** Example of different TCR profiles from low Gini index to high Gini index. The pie chart indicates the fraction of each clone within the repertoire of each sample. The corresponding H&E staining of corresponding FFPE tissue. (**b)** The Gini index distribution histogram of two major types of NSCLC, lung adenocarcinoma (AC) and squamous cell carcinoma (SqCC). (**c)** Boxplot represents the paired comparison of Gini index between tumor and adjacent normal lung tissue (paired Wilcoxon signed-rank test, n = 20). (**d)** Boxplot presents the difference in the Gini index between never-smokers and smokers. (**e)** Kaplan-Meier survival analysis of NSCLC patients stratified based on the Gini index using median cut-off (cut-off = 0.25).

The RNAseq analyses of the 182 NSCLC cases revealed a heterogeneous distribution of samples, in median with lower Gini indexes (median 0.25), with some few exceptions. The distribution of T-cell clonality was similar between adenocarcinoma and squamous cell cancer (**Fig. 1b**). The clonality of 20 corresponding to normal lung tissue from the same patient, revealed generally higher Gini coefficients than tumors (p = 0.021, **fig. 1c**).

### TCR-clonality and association with clinical parameters and survival

Subsequently, we analyzed whether TCR clonality was associated with clinical characteristics. None of the evaluated parameters (patient age, stage, and sex) correlated with the Gini index, with the only exception that ever-smokers revealed a higher TCR receptor clonality in their tumors compared to never-smokers (p = 0.015, **fig. 1d**).

The survival analysis of the Gini index did not show a significant prognostic impact (p = 0.53, median cut-off, **fig. 1e**). This was also true when the Gini index was analyzed as a continuous variable in the univariable Cox regression (HR: 0.89, CI: 0.58–1.36, p=0.59) and multivariable Cox regression model (adjusted to age, stage, and performance status, HR: 0.95 (0.62-1.47), p = 0.83, **table S2**). However, in the descriptive survival analysis without using data censoring at five years and applying an optimal cut-off, a significantly increased survival of patients with higher Gini index, i.e. with dominant clones was found (p = 0.012, **figure S2**).

### The T-cell repertoire in the molecular background of lung cancer

The mutation data of 82 lung cancer-related genes were previously determined by targeted sequencing (*37*) and associated with the Gini index (**Table S3**; **fig. 2a**). We found that the Gini index was higher in tumor harboring p53 mutations (p = 0.001, FDR = 0.023), ARID1A (p = 0.009, FDR=0.18), EPHB6 (p = 0.024, FDR = 0.38), and CSMD3 (p = 0.031, FDR = 0.41).

**Fig. 2.**
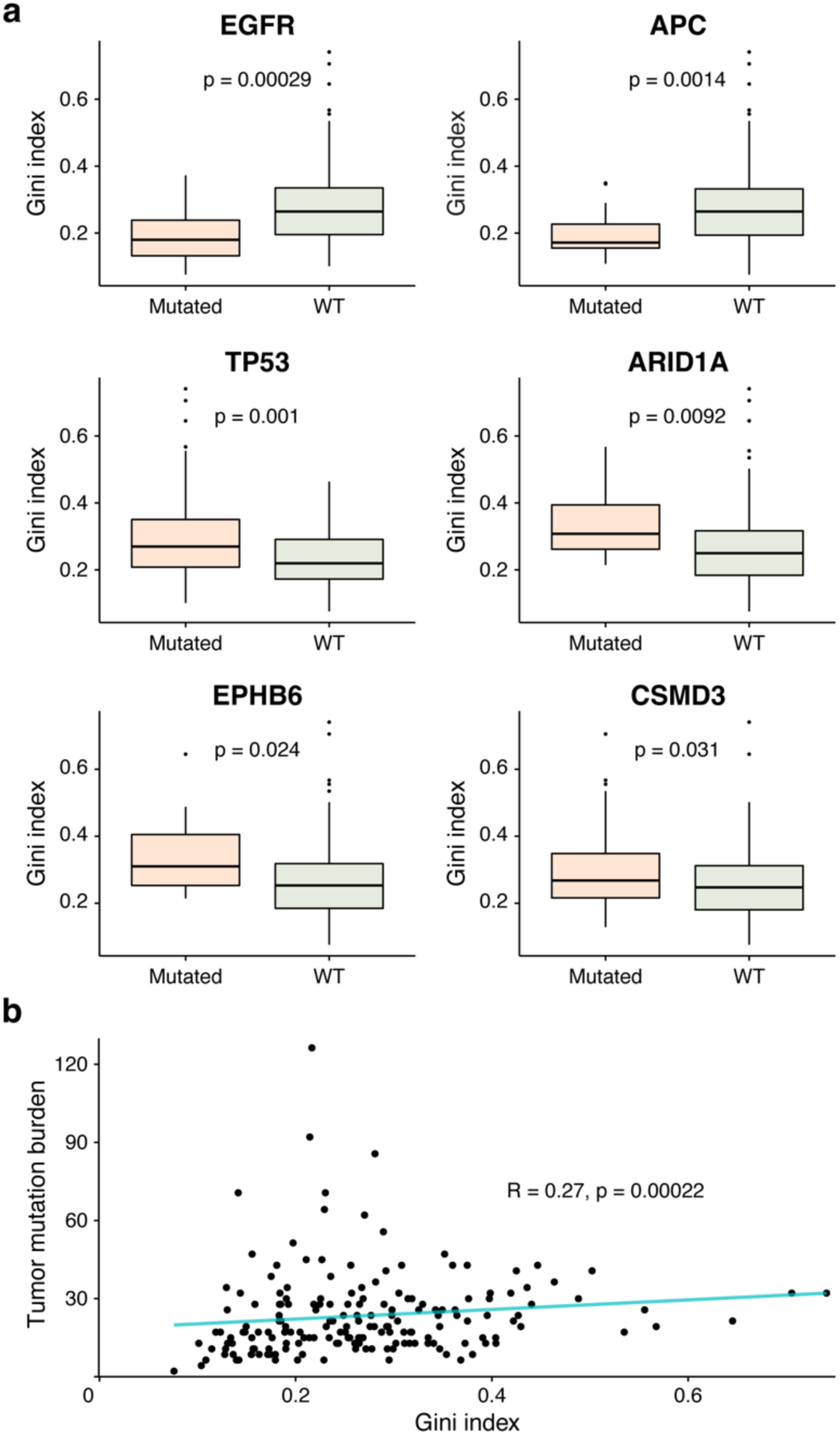
TCR clonality and association with mutations. (**a)** Comparison of Gini index between patients with tumor harboring pathogenic mutations (EGFR, APC, TP53, ARID1A, EPHB6, CSMD3) and tumors without mutations (wildtype, WT). (**b**) Correlation blot between Gini index and estimated tumor mutation burden.

The Gini index was lower for patients with EGFR (p < 0.01, FDR = 0.024) and APC mutations (p < 0.01, FDR = 0.03). Importantly, the Gini index showed a significant positive correlation with the estimated tumor mutation burden (**Fig. 2b**, p < 0.001). Taken together, the genetic background of the tumor and the amount of putative neoantigens have a relation to the TCR clonality in the local tumor environment.

### The T-cell repertoire and the immune landscape of lung cancer

Subsequently, we evaluated how the Gini index was associated with the immune cell repertoire in the microenvironment. When the immune pattern was created by unsupervised clustering (**Fig. 3a**), we observed a higher Gini index in the immune rich (inflamed) cases (**Fig. 3b**, p<0.001). However, the association of high TCR clonality with infiltrating immune cells was pronounced only when the tumor compartment, and not the stromal compartment, was considered (**Fig. 3c**), with a significant correlation to PD-L1 expression on tumor cells (p = 0.003). High Gini index was also strongly associated with CD3 positive lymphocytes (p < 0.001), PD-1 (p < 0.001), CD8 (p < 0.001), CD45RO (p < 0.001), and CD163 positive cells (p < 0.001) in the tumor compartment. By using an established multiplex immune fluorescence pipeline, we were also able to subtype the relevant lymphocyte populations, indicating that predominantly effector CD8 and CD4 cells, and Treg-cells strongly associated with the Gini index (p < 0.001, **Fig. 3d**), although the association of TCR clonality with Treg-cell was restricted to the stroma compartment. In line with the cell density analysis, the distance analysis confirmed that the T-cell clonality is higher when effector CD8 cells are closer to the tumor cells (**Fig. 3e**). Of non-lymphocytic cell types particularly, the number of pro-inflammatory M1 like macrophages correlated with higher T-cell clonality (p < 0.01) (**Fig. 3d**) possibly linking M1 macrophages to antigen presentation.

**Fig. 3.**
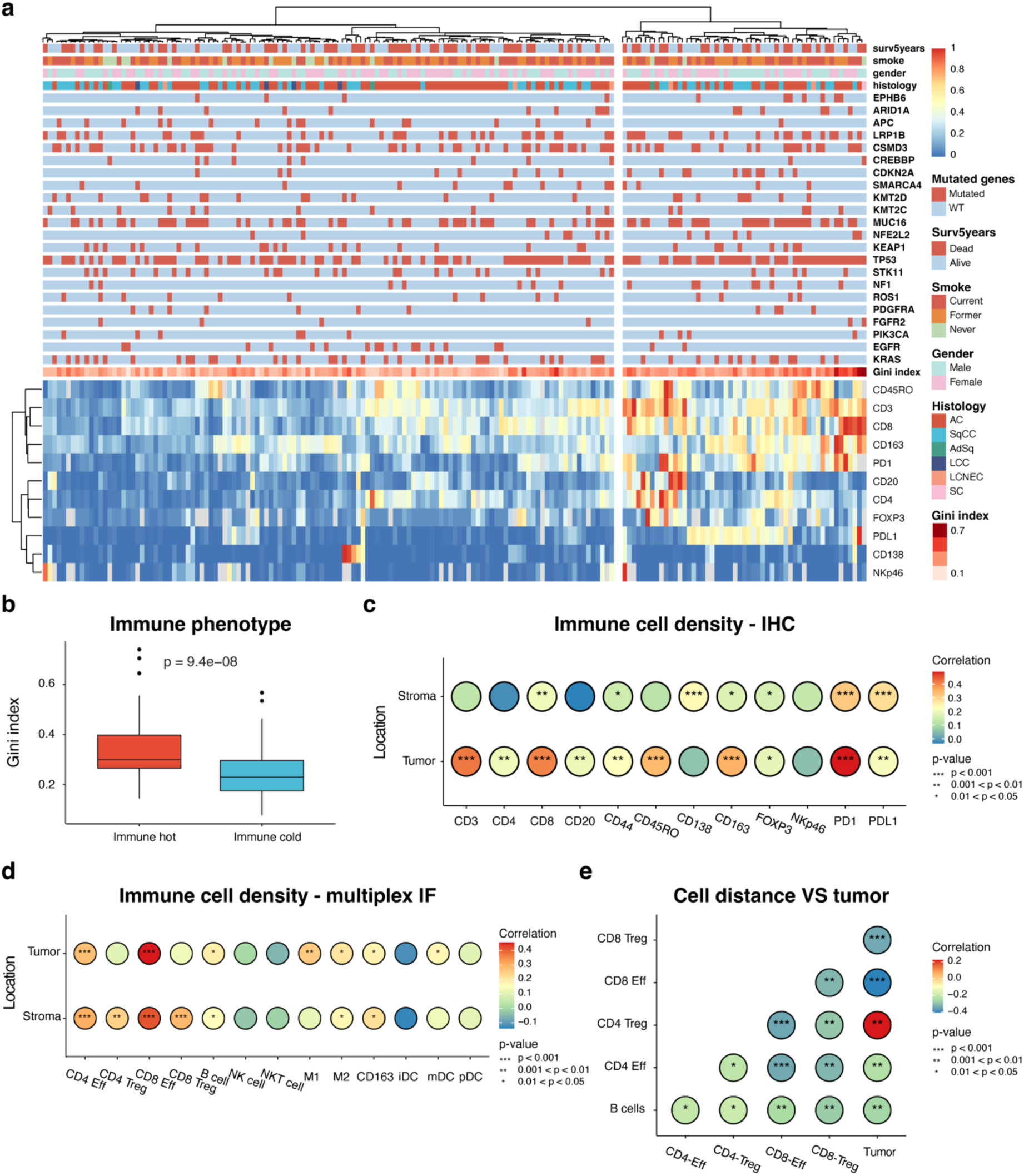
Association of TCR clonality with immunophenotypes. **(a)** Unsupervised hierarchical cluster analysis based on normalized immune markers expression scores assessed using the IHC staining of cancer TMA. Patients were stratified into immune hot and immune cold groups. The 5-year survival (Surv5years), clinical parameters, mutation status for selected genes and their Gini index were given. (**b**) The Gini index of immune hot and cold tumors was compared. (**c**) Immune scores based on immunohistochemical stainings of immune markers were associated with the Gini index for the tumor and stroma compartment separately. The correlation level is indicated by the color. (**d**) Immune cell densities based on multiplex IF were correlated with the Gini index for the tumor and stroma compartment. (**e**) The cell nearest neighbor distances between each immune cell type and tumor cells were correlated with the Gini index. Red color indicates a longer distance between CD4 Treg cells and tumor cells in cases with higher Gini index.

The Gini index is not only related to the absolute numbers of immune cell types, but also correlates with a higher relative proportion of effector CD8 cells in the immune environment (**Fig S3a**), indicating a general change of the immune status.

The observation that local TCR repertoire and the immune phenotypes are interconnected was confirmed in the gene enrichment analysis based on RNAseq data of the top quartile cases with the highest Gini index and the bottom quartile with the lowest Gini index. Higher Gini index demonstrated enrichment of genes related to T-cell activation and expansion, with antigen presentation and binding as well as cytokine signaling. Also, NK cell mediated immunity (*KLRC1*, *KLRC2*, *KLRC3*, *KLRC4*, *NKG7*) was overrepresented in this analysis (**Fig. 4; table S4**). In general, genes that were typically associated with immune activation (*PRF1*, *GZMB*, *GZMA*, *INFG*, *IL17*, *NCR1*) were contrasted by inhibitory signals (*LAG3*, *PD*-*1*, *TIGIT*, *IDO1*) as an indication for T-cell exhaustion (*40–42*).

**Fig. 4.**
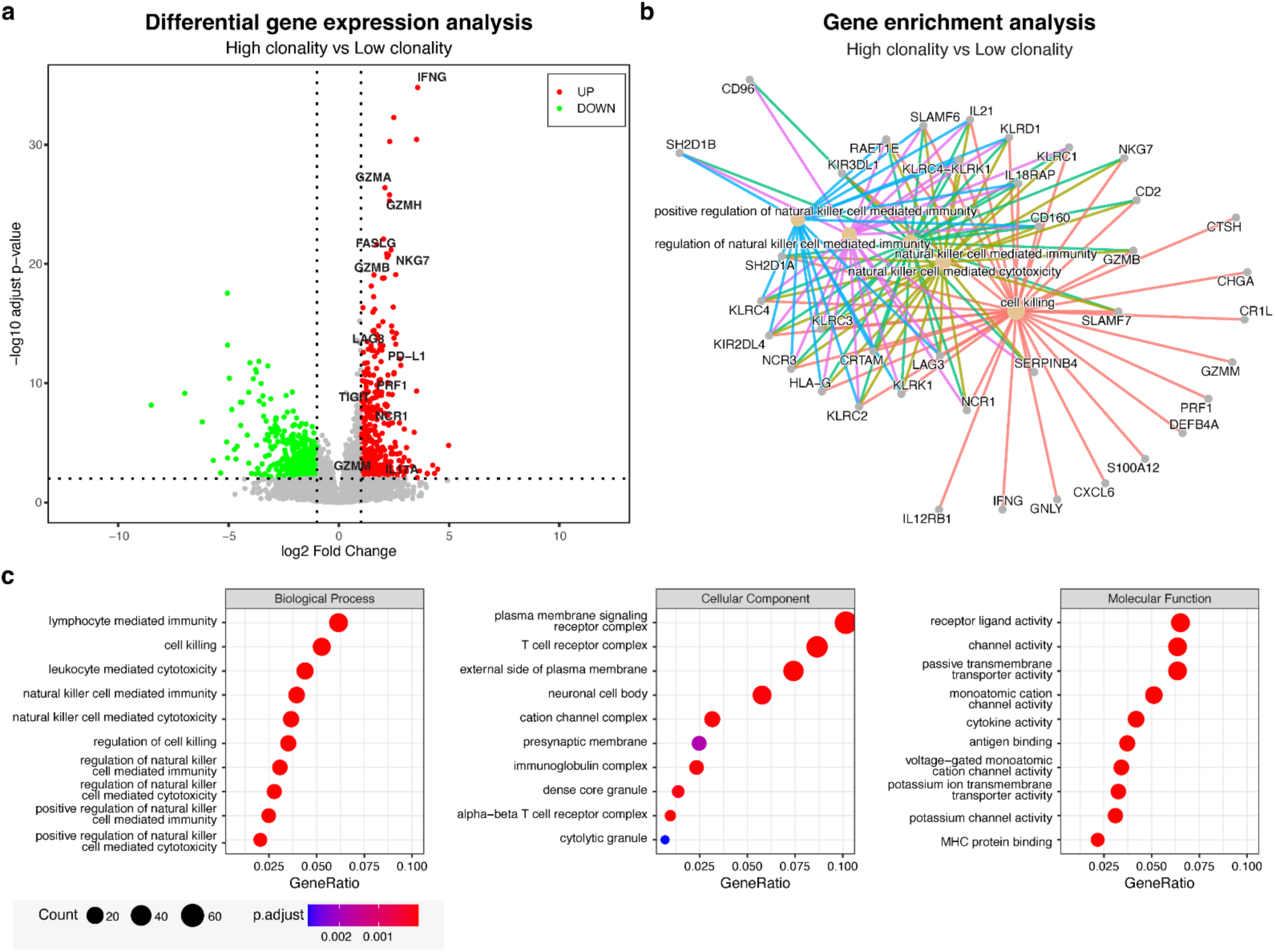
Differential gene expression and gene enrichment analysis. (**a**) Volcano plot indicating the differently expressed gene (adjusted p-value < 0.01) in the top 25% high Gini index tumors compared to 25% cases with the lowest Gini index. The upregulated genes were visualized in red, and downregulated in green. Selected genes were labeled. (**b**) Enrichment plot for connection between differentially expressed genes and their pathway. (**c**) Enrichment plots indicate the biological pathway, cellular component, and molecular function in which differentially expressed genes were involved. Gene ratio (GeneRatio) indicates the percentage of genes involved in the differentially expressed gene set.

Taken together, we confirmed clonality is strongly related to immune activation with signs of T-cell exhaustion, a higher immune cell infiltration with effector cells in the tumor cell compartment.

### Location of dominant T-cell clones in the *in situ* environment of cancer

Recent advances in *in situ* sequencing techniques allow us to precisely localize gene expression in the cancer microenvironment. Here, we use this technique to recover the known variable gene on CDR3 TCR region of a dominant clone in patients’ cancer tissue (**Fig. 5a**). We identified the location of the expanded, dominant T-cell clones as well as minor non-dominant clones. By overlaying corresponding HE staining and tumor cell compartments and stroma cells compartments were annotated based on *EPCAM*, *CDH1* mRNA expression (**Figure 5b**). Based on spatial binning, dominant TCR clones were more abundant within the tumor cell compartment in all three analyzed patient samples **(Fig. 5c)**, indicating that expanded TCR clones are more frequent in direct contact with cancer cells. These expanded T-cell clones were more often of CD8 subtype, confirming the CD8 expansion observed in the bulk RNA sequencing data (**Fig. S3b, S3c)**.

**Fig. 5.**
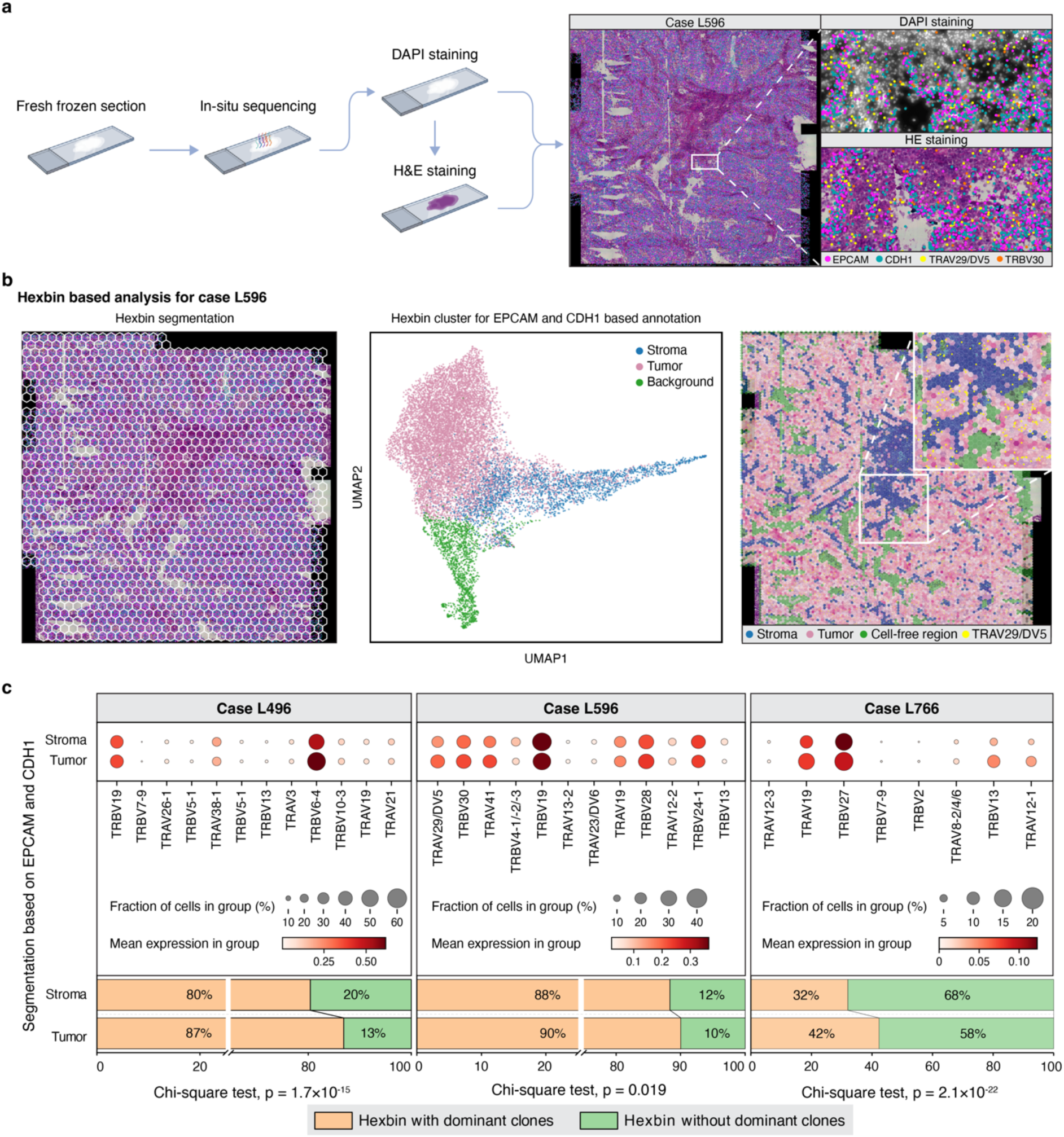
Hexbin based analysis of *in situ* sequencing data. (**a**) *In situ* sequencing (ISS) workflow, illustrated by case L596. The right panel shows stained image with markers for EPCAM, CDH1, as well as the TCR genes TRAV29/DV5 and TRBV30). (**b**) Hexbin-based analysis pipeline applied to the ISS data illustrated by case L596. The left image depicts the hexagonal segmentation of the ISS results, where each hexbin aggregates detected signals based on their spatial coordinates. The middle UMAP plot visualizes the clustering of all marker genes, with the tissue compartments (tumor versus stroma) annotated based on the expression levels of EPCAM and CDH1. UMAP clusters were used to compartmentalize the tissue section (right figure) into tumor, stroma, and background, along with the location of a dominant clone. (**c**) Comparison of dominant TCR clones between tumor and stroma compartments across three cases (L496, L596, L766). The dot plot illustrates the proportion of hexbins containing each dominant clone in tumor and stroma: The dot size indicates the fraction of hexbin containing the dominant clone. The color intensity indicates the mean expression of the dominant clone transcripts. Chi-square tests were conducted to assess the distribution of dominant clones between tumor and stroma, with p-values indicated for each case.

### T-cell clonality and the response to Immune checkpoint inhibitors

Cancer cells might evade immune surveillance by the expression of checkpoint molecules like PD-L1 on cancer cells. Thus, checkpoint inhibitor treatment should be particularly effective in tumors with expanded T-cell clones i.e., a high Gini index.

To test this hypothesis, we evaluated an immune checkpoint-treated cohort of NSCLC patients (n = 9) with publicly available RNAseq data (*33*). The Gini index was significantly higher in patients who responded to checkpoint inhibitor treatment (**Fig. 6a**). This association was also seen in the survival analysis of these advanced NSCLC patients, with a difference in median survival of 13.8 compared to 2.9 months (log-rank test; p = 0.0027, **fig. 6b**). Similarly to the Uppsala cohort, differential gene expression analysis of this mRNA data set indicated that TCR clonality was associated with marker of T-cell activation (*GZMA*, *PRF1*, *IL2RB*) and exhaustion (*CD244*, *CD96*); and also involves NK cell immunity (*KRLB1*, *KLRK1*, *KKG7*) **(Table S5).**

**Fig. 6.**
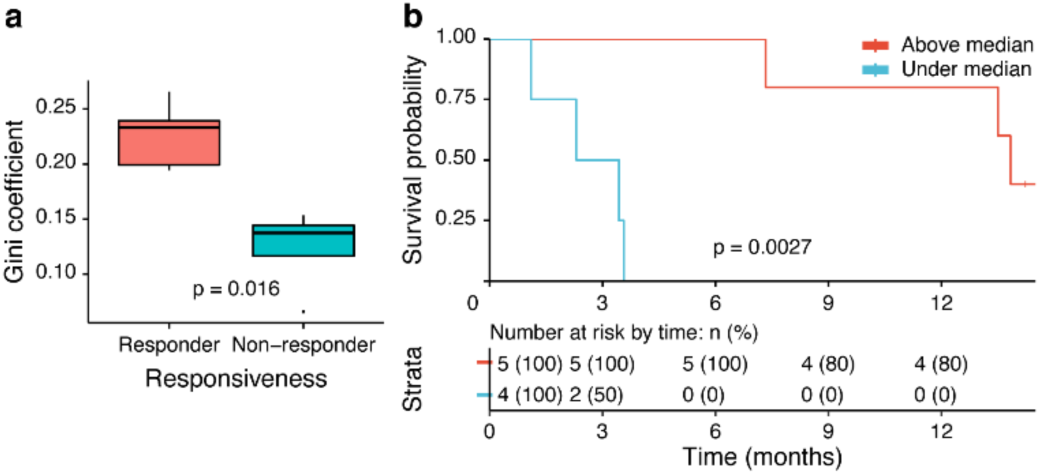
TCR clonality in NSCLC patients treated with checkpoint inhibitor therapy. (**a**) Box plot for comparison of Gini index between NSCLC patients who responded (n=5; partial response and complete response) to immunotherapy and non-responders (n = 4; stable disease and progressive disease). (n=9) (**b**) Kaplan-Meier survival analysis of the patients stratified by median Gini index cut-off (cut-off = 0.19)

## Discussion

Our study uncovers a significant variability of T-cell clonal expansion in human lung cancer tissue. The expansion, quantified as Gini index, is associated with a specific molecular background (e.g. EGFR, TP53, APC), higher mutational load, and activated but also exhausted immune cell profiles. A prominent potential clinical relevance of the findings is demonstrated, as patients with elevated TCR clonality benefit from checkpoint inhibitor therapies.

Our study is based on previous developments of bioinformatic pipelines, allowing for the precise alignment of TCR sequences from crude RNA sequencing data to estimate the number of T-cell clones (*26, 31*). Our bioinformatic approach was adopted from the study of Valpione (*26*) and Farmanbar (*31*) who analyzed TCR clonality in melanoma and lymphoma respectively.

In our study, we used the Gini index instead of entropy to describe the distribution of TCR clones, which emphasize unevenness and might be more sensitive to identifying clonal expansion (*17*).

Our method was further validated against an independent CE-marked, DNA-based assay (Lymphotrack), which is used in clinical diagnostics to identify and track clonality in lymphocytic diseases (*20*). The strong congruence confirmed the robustness of our pipeline and the accuracy of our TCR data set.

There are few previous studies analyzing TCR clonality in diagnostic lung cancer samples. The study of Zhang (*43*) evaluated tumor tissue of 10 patients who received neoadjuvant immunotherapy. They found that the entropy of TCR clones in the remaining tumor bed after immunotherapy correlated with the residual tumor cell viability and major pathological response. They also showed, using pre- and posttreatment blood samples, an expansion of corresponding dominant clones. In a comparable approach using a targeted RNA assay, Casarrubios and coworkers (*44*) confirmed those results in post-treatment tumor tissue. In addition, they found in the pre-treatment tissue that lower TCR evenness and higher proportion of top 1% T-cell clones, were associated with response to neoadjuvant therapy.

A more recent study from the group of Amos (*29*) compared complex immune profiles in tumor and normal lung tissue from surgical specimens of 67 NSCLC patients. Notably, their TCR clonality analysis identified substantially fewer T-cell clones (total number of reads), although they used the same MXCR pipeline. This discrepancy might be due to a different quality of RNAseq source data. However, analogous to our study they found a higher TCR clonality in the normal tissue. A larger study analyzed TCR clonality in blood and tissue samples (tumor and adjacent normal lung) of 214 patients with a targeted RNA assay. Their findings indicated a strong relation between TCR entropy and specific immune cell profiles. When filtrating tumor TCR against blood clonality they demonstrated that higher “tumor enriched” clonal expansion in the normal lung was associated with longer survival in patient group after surgical resection (*22*).

Our study confirmed and extended previous findings in an independent, molecularly and clinically extensively annotated data set of NSCLC patients. It should be stressed that the frequency and richness of detected TCR clones in our RNAseq data were exceedingly higher than in all previous studies, and we also validated our analysis pipeline. Our comparative analysis provided evidence that the clonal expansion of T-cells is paralleled by the infiltration of CD8 effector cells in the tumor cell compartments, i.e. they are in direct contact with tumor cells. We confirmed this relation in the spatial multiplex IF analyses and even visualized these expanded T-cell clones in the *in situ* cancer environment by *in situ* RNA sequencing. This *in situ* data positioned the expanded TCR clones closer to the tumor cells and confirmed that they are rather of CD8 subtype.

Together, the *in situ* findings strongly indicate that some expanded TCR clones, are functionally related to tumor antigens. Therefore, it was not surprising that the potentially specific immune response coincided with an increase in inhibitory cell populations (CD163 M2-like macrophages) and markers of immune cell exhaustion (LAG3, PD-L1, PD1), possibly counteracting any beneficial cytolytic response.

The observation that T-cell expansion was associated with an ineffective immune reaction might explain why higher TCR clonality in tumor tissue does not translate into a survival benefit in our surgically treated patient cohort and possibly also in the surgical-treated cohort previously described by Reuben (*22*). In contrast, when we analyzed the NSCLC cohort of advanced patients that received checkpoint inhibitor therapy, we found a significant association between TCR clonality to response to therapy and overall survival, although the cohort was fairly small.

Taken together our findings support the concept that the clonal expansion is tumor-antigen specific but ineffective and can thus potentially be therapeutically unleashed. This was in line with previous studies of neoadjuvant-treated patients, where TCR clonality was related to pathological response in the post-treatment surgical specimens (*43, 44*).

Nevertheless, a possible explanation is, that the TCR clonality is only a strong surrogate for the immune environment and hence not related to a specific pretherapeutic anti-tumor immune reaction. Although our Uppsala cohort indeed presents a large, highly detailed TCR clonality data set, the immunotherapy cohort consisted only of 9 patients, with limited statistical power. The low number reflects the problem that fresh tissue is usually not available from advanced cancer patients but is still required to obtain high-quality RNA for the TCR clonality analysis. This limitation hinders further evaluation of the TCR clonality, based on RNAseq, as a predictive biomarker for immune therapy. As an alternative, a more robust targeted DNA-based assay might provide similar information as the RNAseq data and would be applicable to minute FFPE material in clinical diagnostics.

In conclusion, our study linked T-cell clonal expansion to relevant molecular and immunophenotypes of NSCLC and provided evidence that the analysis has potential as a predictive marker delineating a group of patients that will benefit from checkpoint inhibition. Our publicly available, extensive data set also presents a unique source for more focused studies, aiming to understand the mechanism of immune cell activation in lung cancer. Hopefully, this will contribute to improving the current therapeutic options with only limited overall response.

### Declaration of generative AI and AI-assisted technologies in the writing process

During the preparation of this work the authors used ChatGPT 4.0 (Open AI) for language editing. After using this tool the authors reviewed and edited the content as needed and take full responsibility for the content of the published article.

## Acknowledgment

This study was partly supported by the Sjöberg Foundation, Sweden; the Swedish Cancer Society, the Lions Cancer Foundation Uppsala, Sweden, and the Swedish Government Grant for Clinical Research. CS hold a starting grant from the Trond Mohn Foundation. We thank the Research and Development Unit of the Pathology Department of the University Hospital Uppsala for their continuous support as well as the Uppsala Biobank.

## Competing interests

All authors declare no conflicts of interest.

---

**Fig. S1.**
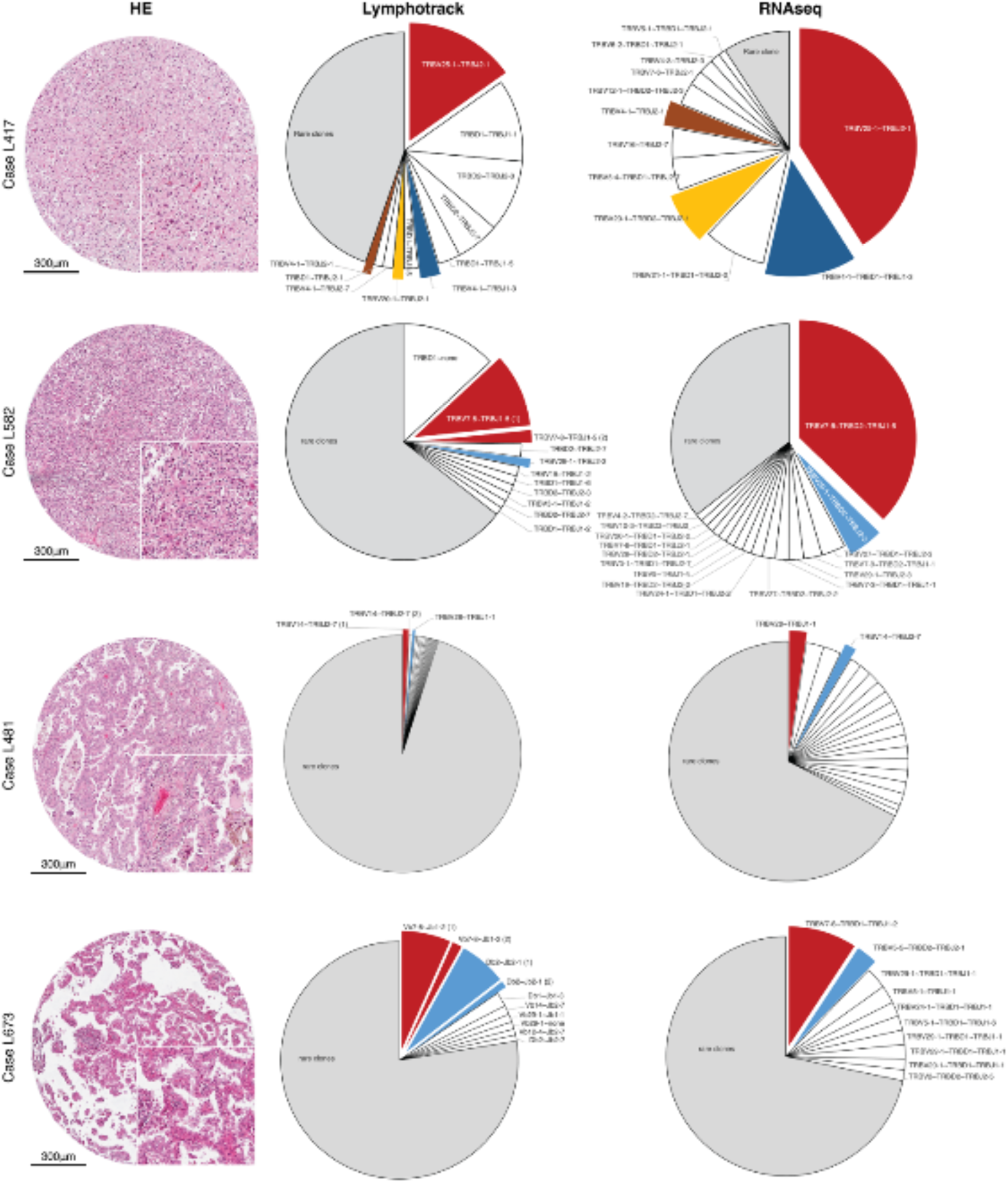
Validation of TCR clonality analysis. The Lymphotrack assay was used to verify our RNAseq based pipeline to determine the distribution of TCR clones. Each row presents one case with corresponding H&E staining of the cancer tissue. The pie charts indicate the identified T-cell clones with Lymphotrack and RNAseq. The same clones are visualized with the same color.

**Fig. S2.**
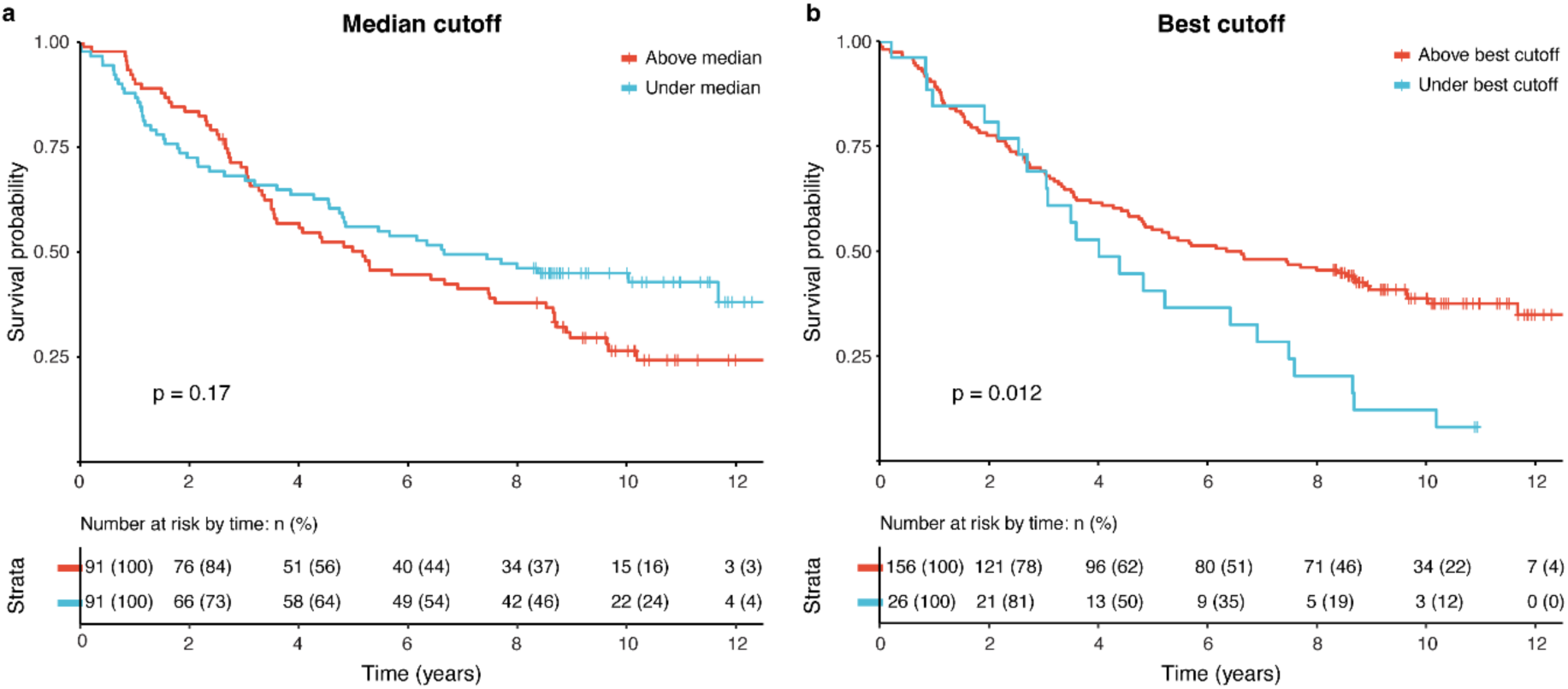
TCR clonality and NSCLC patient survival. (**a**) Kaplan-Meier overall survival analysis of NSCLC patients without time censoring stratified based on the Gini index using median cut-off (cut-off = 0.25). (**b**) Kaplan-Meier survival analysis of NSCLC patients without time censoring stratified based on the Gini index with the best cut-off calculation (cut-off = 0.16)

**Fig. S3.**
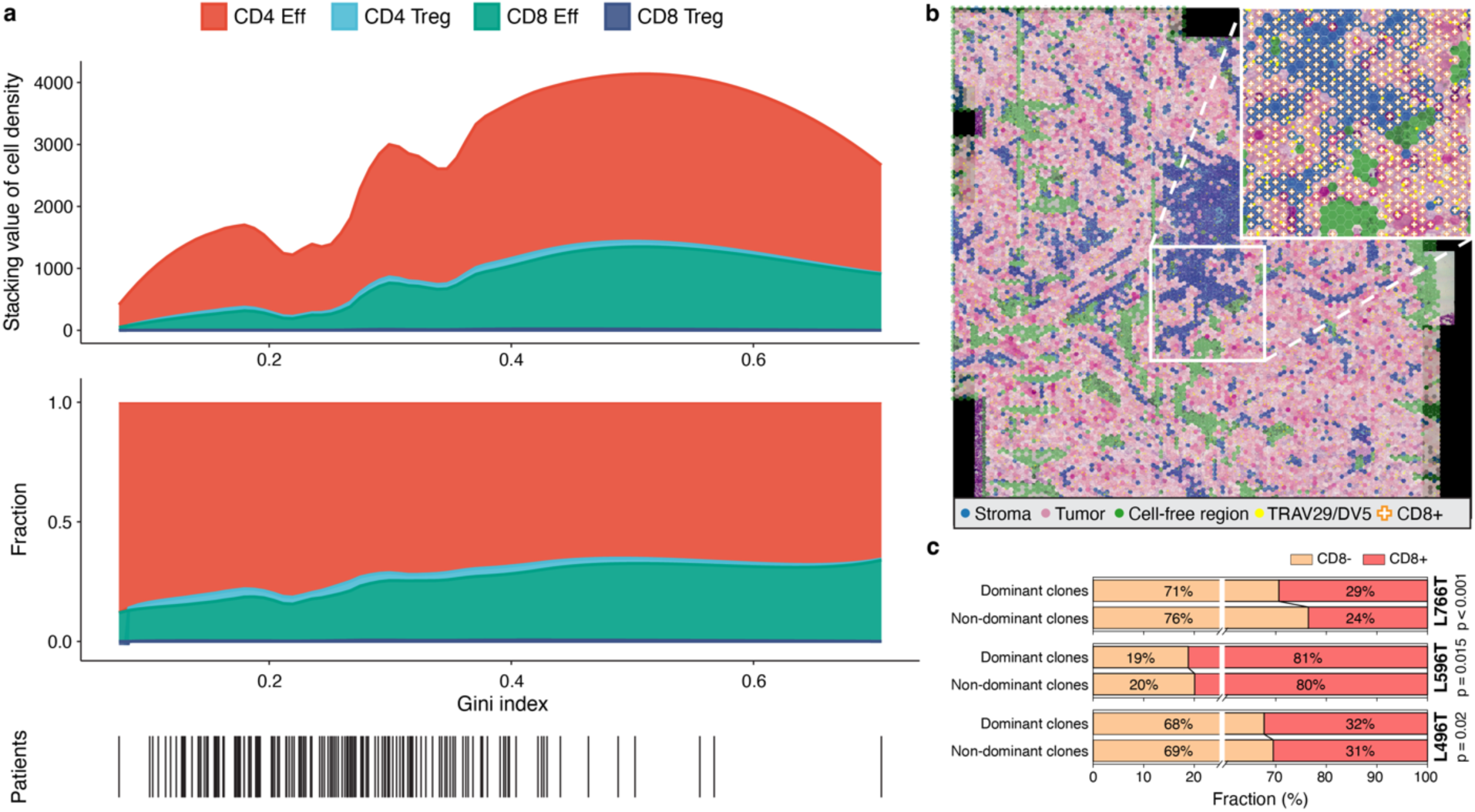
TCR clonality and association with T cell phenotypes. (**a**) Change of T-lymphocyte composition in relation to Gini index. The 4 major T cell subtypes were quantified by multiplex IF and plotted against the Gini index: CD4 effector cells (CD4 Eff), CD4 regulatory cells (CD4 Treg), CD8 effector cells (CD8 Eff) and CD8 regulatory cells (CD8 Treg). The upper figure shows the absolute change in cell densities; the lower figure shows the relative fraction of T-cell subsets. Below the patient distribution is indicated. (**b**) *In situ* map based on hexagon bins indicating tissue regions and the presence of the dominant clone TRV29/DV5 and CD8 cells. (**c**) Comparison of CD8 positive bins containing dominant and non-dominant clones.

**Table S2:** Cox regression survival analysis of NSCLC patients.

| Cox regression |  |  |
| --- | --- | --- |
| Univariable analysis |  |  |
|  | HR (95% CI) | p-value |
| Gini index (Above median / Under median) | 0.89 (0.58 - 1.36) | 0.59 |
| Multivariable analysis |  |  |
|  | HR (95% CI) | p-value |
| Gini index (Above median / Under median) | 0.95 (0.62 - 1.47) | 0.83 |
| Age (Above 70 / Under 70) | 1.26 (0.82 - 1.93) | 0.29 |
| Gender ( Female / Male) | 0.78 (0.51 - 1.21) | 0.27 |
| Smoke (Current&Former / Never) | 1.27 (0.59 - 2.72) | 0.55 |
| Histology (AC as reference) |  |  |
| SqCC | 0.83 (0.52 - 1.33) | 0.43 |
| Other | 0.82 (0.34 - 1.95) | 0.64 |
| Stage (II-IV / I) | 1.80 (1.16 - 2.77) | 0.0087 |

## Notes

### Competing Interest Statement

The authors have declared no competing interest.

